# Dynamic regulation of NeuroD1 expression level by a novel viral construct during astrocyte-to-neuron reprogramming

**DOI:** 10.1101/2025.02.17.638625

**Authors:** Natalie Mseis-Jackson, Mei Jiang, Mehek Sharma, Arya Ranchod, Christine Williams, Xuanyu Chen, Hedong Li

## Abstract

Astrocyte-to-neuron reprogramming presents a viable approach for regenerative medicine. The reprogramming factor NeuroD1 has demonstrated capability of neuronal reprogramming with high efficiency both in culture and in the injured central nervous system. High level of NeuroD1 expression is required to break down the cellular identity barrier for a successful reprogramming, and yet persistence of this high level drives the reprogrammed neurons primarily to glutamatergic subtype. This is consistent with the critical role of NeuroD1 in determination of glutamatergic neuronal lineage during development. However, diversified neuronal subtypes are needed to establish appropriate neuronal connectivity in disease/injury conditions. We reason that continuously high level of NeuroD1 expression forces the reprogrammed neurons into glutamatergic subtype, and that reducing NeuroD1 level after reprogramming may allow generation of neurons with diversified subtypes. For this purpose, we engineered a novel viral expression vector by which NeuroD1 expression can be dynamically regulated during the reprogramming process. Specifically, the target site of a neuron-specific microRNA (miR-124) is incorporated in the expression system. Therefore, this novel construct would still achieve a high NeuroD1 expression level in astrocytes for reprogramming to occur and yet reduce its level in the reprogrammed neurons by suppression of endogenous miR-124. In this study, we demonstrated that this construct elicits a dynamic gene expression pattern with much reduced level of NeuroD1 at later stages of neuronal reprogramming. We also showed that this construct still retains relatively high reprogramming efficiency and can generate mature neurons with an enhanced GABAergic neuronal phenotype.

## Introduction

In vivo neuronal reprogramming has recently emerged as a novel technology to regenerate new neurons from endogenous glial cells in the central nervous system (CNS) simply by overexpressing certain neurogenic transcription factors^1,2^. Since no foreign cells are involved, this regenerative approach bypasses the critical hurdle of immunorejection that cell transplantation therapy is facing. NeuroD1 is such a neurogenic transcription factor that can reprogram endogenous glial cells into functional neurons in the injured brain^3^ and spinal cord^4^. NeuroD1-mediated neuronal reprogramming has been demonstrated in different disease/injury models despite an existing debate on the validation and efficiency of neuronal reprogramming^5,6^. One interesting observation is that NeuroD1-reprogrammed neurons from astrocytes are mostly glutamatergic (i.e. excitatory) subtype^3^. This is likely because NeuroD1 is one of the critical determination factors that specify glutamatergic neuron-lineage during brain development^7,8^. However, diversified neuronal subtypes, i.e. both excitatory (E) and inhibitory (I) neurons, would be needed to keep the E/I balance in the reconstructed neuronal circuitry for functional repair after disease/injury. Although other transcription factors such as Ascl1/Mash1 and Dlx2 can reprogram astrocytes into GABAergic neurons, their neuronal reprogramming efficiency is much lower than that of NeuroD1^9,10^. Therefore, a tempting strategy is to modulate NeuroD1-mediated neuronal reprogramming so that diversified neuronal subtypes may be generated while still maintaining high reprogramming efficiency.

Interestingly, NeuroD1 has been shown to compete with Mash1/Ascl1 in determination of neuronal subtype during forebrain development^11^, suggesting that relative gene expression levels between these transcription factors may be crucial for neuronal subtype output. For reprogramming to occur, NeuroD1 expression must reach a high enough level^12,13^. However, consecutively high expression of NeuroD1 by the strong promoters such as CAG may force the reprogrammed neurons into glutamatergic subtype since NeuroD1-positive neurons in the adult brain are all glutamatergic^14^. Therefore, we set out to determine if we can generate inhibitory neurons using NeuroD1 by reducing its expression level after neuronal reprogramming has occurred. Toward that, we have engineered a novel viral expression vector by which NeuroD1 expression can be dynamically regulated during the reprogramming process.

MicroRNAs (miRNAs) are endogenously derived, short non-coding RNAs that regulate gene expression in a post-transcriptional manner^15^. In collaborating with transcriptional controls by transcription factors, miRNAs play important roles in many, if not all, biological processes including differentiation of neural cell types during CNS development^16^ and pathological processes under disease/injury conditions^17–22^. Our previous work also demonstrated that miRNAs are critical to cellular differentiation in the developing mouse forebrain^23^ and cerebellum^24,25^, and indispensable to reactive astrogliosis after spinal cord injury (SCI)^26^. MiRNAs generally bind to the 3’-untranslated region (UTR) of their target mRNAs through imperfect base-pairing and perform posttranscriptional regulation on gene expression by either degrading the mRNA targets or inhibiting their translation^15,18,27–31^. Based on these mechanisms, vectors have been designed to improve cell type-specificity of gene expression using the silencing effect of miRNAs in combination of cell-type specific promoters. For example, miR-124 is expressed specifically in neurons but not astrocytes^32^. Insertion of a miR-124 target sequence at the 3’-end of a transgene in an expression vector results in post-transcriptional silencing of the transgene in neurons but not astrocytes^33–37^, due to the neuron-specific expression of miR-124^13^. In a recent report, applying target sequences of multiple neuron-specific miRNAs including miR-124 can further improve astrocytic specificity of gene expression^38^.

In our current study, we integrate this miR-124-mediated silencing technique into our NeuroD1 expression vector driven by a strong promoter CAG^3^. Therefore, during astrocyte-to-neuron reprogramming, high level of NeuroD1 expression in astrocytes (low in miR-124 level) can be achieved for neuronal reprogramming to occur, and then expression level of NeuroD1 is reduced in the reprogrammed neurons (high in miR-124 level), which may allow generation of inhibitory neuronal subtypes. To test this idea, we generated a novel NeuroD1 expression construct containing miR-124 target sequences. We showed that this novel construct retains relatively high astrocyte-to-neuron reprogramming efficiency and elicits dynamically regulated NeuroD1 expression levels during reprogramming. More importantly, reduced NeuroD1 expression level in the reprogrammed neurons by this construct weakens the glutamatergic neuronal markers while enhancing the GABAergic neuronal markers.

## Materials and Methods

### Construction of retroviral vectors and viral particle packaging

Several retroviral expression constructs were used in this study. Most of these constructs share the same vector backbone that contains a CAG universal promoter to drive high levels of gene expression. *Retro-GFP*, *Retro-ND1-GFP* and *Retro-ND1-RFP* were reported previously^3,39^ while *Retro-ND1-124T-GFP* and *Retro-124T-GFP* were newly constructed for this study. Specifically, to create miR-124 target sites (*miR-124T*), we modified a reported strategy to use the perfectly complimentary sequence of mature miR-124 (*TGGCATTCACCGCGTGCCTTA*) instead of the natural target sequence of miR-124 from the *integrin-β1* gene^33^. This is intended to maximize miRNA-mediated repressive effect on gene expression^40–42^. Similarly, we also inserted a single adenosine residue to improve sensitivity to miR-124 as demonstrated^43^. Eight copies of *miR-124T* were designed with a *HindIII* site in the middle (4x miR-124T-*HindIII*-4x miR-124T). The sequence is *TGGCATTCACCGCGTGCCTTAgagaTGGCATTCACCGCGTGCCTTAgagaTGGCATTCACCGCGTGCCTTAg agaTGGCATTCACCGCGTGCCTTAaagcttTGGCATTCACCGCGTGCCTTAgagaTGGCATTCACCGCGTGCCTTAgagaTGGCATTCACCGCGTGCCTTAgagaTGGCATTCACCGCGTGCCTTA*. The 8x *miR-124T* was synthesized as oligonucleotides and subsequently cloned into Retrovirus expression vectors using PCR-based techniques. For *Retro-ND1-124T-GFP*, the 8x *miR-124T* was inserted into *Retro-ND1-GFP* between *ND1* coding sequence and *IRES* by using *PmeI* and *NotI* sites. For *Retro-124T-GFP*, the 8x *miR-124T* was inserted into *Retro-ND1-GFP* to replace *ND1* coding sequence by using *SfiI* and *PmeI* sites. Lastly, we also generated a miR-124 overexpression construct using a MSCV based expression vector. Briefly, a 496-bp genomic fragment containing miR-124^44^ was subcloned into *pmCherry-miR-125b-1*^45^ (a gift from David Baltimore, Addgene plasmid #58990) using *NotI* and *XhoI* sites to replace *miR-125b-1*. All new constructs were sequenced to confirm the correctness of the inserted DNA fragments.

Viral particle packaging was performed as described^39^. For miRNA overexpression vector packaging, a plasmid containing human Dicer shRNA [*pSicoR human Dicer1*, a gift from Tyler Jacks (Addgene plasmid #14763)] was included to increase virus yield ^46,47^. Virus-containing culture media were centrifuged and filtered to remove cell debris, then aliquoted and stored at -80°C before use. Virus media were routinely assayed for titer by infecting HEK293T cells. Our retrovirus titers are usually at 10^7^ genomic copies (GC)/ml.

### Cell culturing, plasmid transfection, and retroviral infection

The procedures for human astrocyte (HA) cultures (HA1800, ScienCell) and medium change after virus infection were previously described^39^. For retroviral infection experiments, spinfection was performed to increase infection efficiency^48^. Briefly, virus-containing media were mixed with polybrene (4 µg/ml) before being applied onto HA cultures. The resulting culture plates were centrifuged at 1,000 × g for 45 min at room temperature and then cultured in CO_2_ incubator at 37°C. At one day post virus infection, cultures were subject to medium change from HA culture medium (HAM) to neuron differentiation medium (NDM)^39^. For long-term cultures, NDM was changed every 2-4 days. Brain-derived neurotrophic factor (BDNF, 20 ng/ml, Invitrogen) was supplemented to support long-term survival of reprogrammed neurons^39^.

We cultured HeLa cells in Dulbecco’s Modified Eagle Medium (DMEM) supplemented with 10% fetal bovine serum (FBS) and 1% penicillin/streptomycin (P/S). For retroviral infection experiments on HeLa cells, the procedures were similar to those for HA cells except for no medium change after viral infection. For long-term viral infection experiments, infected Hela cells were grown in medium containing 2% FBS after infection to prevent overgrowth^49^. For plasmid transfection experiments, HeLa cell cultures with ∼60% confluency were transfected with total of 2ug DNA per well on a 12-well plate by PEI protocol^39^. The *ND1-GFP* and *ND1-124T-GFP* plasmids were co-transfected with either a control *mCherry* plasmid or the *miR-124-mCherry* plasmid in 1:3 ratio. The HeLa cell cultures were then harvested for Western blot analysis 3 days after transfection.

### HA spheroid formation and culturing

Retrovirally infected HA cultures were passaged 1 day post infection and subject to cellular spheroid formation in a hanging drop fashion following a published protocol^50^. Briefly, ∼10^5^ cells per 20 µl HAM medium were dropped on the lid of culture dishes and inverted to form hanging drops before putting into CO_2_ incubator. The next day, cellular spheroids were formed at the bottom of the hanging drops and transferred onto coverslips with a monolayer of growing HA cells. The HA spheroids were usually attached to the coverslips within 1-2 days after plating. Once attached, HAM was replaced with NDM to facilitate neuronal reprogramming. For long-term cultures, NDM was changed every 2-4 days until cells were fixed for immunostaining. BDNF (20 ng/ml, Invitrogen) was supplemented to support long-term survival of reprogrammed neurons^39^.

### Immunocytochemistry

Immunocytochemistry was carried out as previously described^39^. Briefly, fixed cell cultures were incubated with monoclonal antibodies against NeuroD1 (mouse IgG, 1:1000, Abnova), GFP (rat IgG2a, 1:400, BioLegend), HuD/ELAVL4 (mouse IgG, 1:200, Santa Cruz), GAD67 (mouse IgG, 1:200, Millipore), and NeuN (mouse IgG, 1:400, Millipore); polyclonal antibodies against GFP (chicken IgY, 1:400, Aves), mCherry/RFP (rabbit IgG, 1:500, Abcam), mCherry/RFP (chicken IgY, 1:400, Aves), Map2 (rabbit IgG, 1:400, Abcam), DCX (rabbit IgG, 1:500, Abcam), vGlut1 (guinea pig IgG, 1:200, Millipore), Ctip2 (guinea pig IgG, 1:200, Synapse Systems), anti-SV2 (rabbit, 1:2000, Developmental Studies Hybridoma Bank), GABA (rabbit IgG, 1:200, Calbiochem), and DCX (guinea pig IgG, 1:1000, Millipore), followed by appropriate species-specific secondary antibodies (Molecular Probes). DAPI (10 µg/ml, Sigma) was often included in the secondary antibody incubations to label nuclei. The stained cells were then mounted in mounting medium and analyzed by conventional or confocal fluorescence microscopy.

### Western blot

Western blot analysis was performed as described^39^. Briefly, cell cultures were harvested in RIPA buffer (Alfa Aesar) following manufacturer’s instructions. Forty µg of protein were boiled in SDS sample buffer for 5 min and loaded on each lane of Any kD™ Mini-PROTEAN® TGX™ precast polyacrylamide gels and transferred onto PVDF membranes. The primary antibodies were anti-NeuroD1 (mouse IgG, 1:1000, Abnova) and anti-GFP (rabbit IgG, 1:500, Abcam). The primary antibodies were detected by appropriate species-specific DyLight 700 or 800-conjugated secondary antibodies (1:10,000, Thermo Scientific). Anti-GAPDH (mouse IgG, 1:1000, Sigma) was used to normalize sample loadings. Quantification of relative protein expression levels was done by measuring signal intensity of target bands on a LI-COR Odyssey® Infrared Imaging System and normalizing to that of GAPDH.

### Fluorescence activated cell sorting (FACS) analysis

Both HA and Hela cells were infected with ND1-GFP and ND1-124T-GFP retroviruses separately. At different timepoints (days) post infection, cell cultures were passaged and resuspended in medium containing 2% FBS. The single cell suspensions were kept on ice before analysis. To exclude non-viable cells, 1 μl of 7-AAD staining solution (Enzo, 1 mg/ml) was added to cell suspension prior to FACS analysis on the Agilent NovoCyte Quanteon Flow Cytometer (Agilent Technologies). A gating strategy was applied to exclude cellular debris and small cell aggregates. Only live single cells were analyzed for their GFP expression. Uninfected cells (GFP-) were used as negative controls. There was good separation between GFP- and GFP+ cells in each sample based on their GFP intensity. The GFP median intensity was collected from GFP+ population for each sample and plotted in time course.

### Measurements and statistical analysis

The levels of cellular fluorescence from fluorescence microscopy images were determined in ImageJ software and corrected to mean fluorescence of background readings as previously described^39^. In some experiments, a “circling” method was implemented. Briefly, instead of outlining the entire cell body with tracing on the fluorescent images^39^, a circle was placed within the cytoplasm around the nucleus of each cell, and the fluorescence intensity was measured within the circle. We found that the “circling” method gave similar comparison results to our previous method^39^ and greatly simplified the procedure. After data collection from ImageJ, the corrected cellular fluorescence intensities were then normalized by the average intensity of the *ND1-GFP* group within each experiment before statistical analysis. For quantifications on western blots, data were collected from at least three biological replicates. The data are presented as mean ± SEM. Statistical analysis was performed in GraphPad Prism 9 using Student’s *t*-test. P < 0.05 was considered a significant difference.

## Results

### Design and validation of a novel NeuroD1 expression construct containing the miR-124 target sequence

Based on published information^33^, we synthesized a long oligonucleotide containing 8 copies of the miR-124 target sequence that is completely complimentary to mature miR-124 sequence (**Fig. 1A**) and cloned it at the 3’-end of the NeuroD1-coding sequence of the original vector to generate the novel NeuroD1 expression construct (*ND1-124T-GFP*) for this study (**Fig. 1B**). For the need of our analyses, we have also generated a construct containing miR-124 target sequences alone (*124T-GFP*) and a miR-124 overexpression construct (*miR-124-mCherry*), and included the original NeuroD1 expression vectors with either GFP or RFP as a reporter (**Fig. 1B**). To validate the responsiveness of the new construct *ND1-124T-GFP* to miR-124 level, we performed a co-transfection experiment in HeLa cells followed by western blot analysis. The results showed that, while overexpression of miR-124 had no effect on NeuroD1 protein level by the original *ND1-GFP* construct (**Fig. 1C**), it significantly reduced NeuroD1 protein level by the new construct *ND1-124T-GFP* (**Fig. 1D**). Therefore, the *ND1-124T-GFP* construct can respond to miR-124 level and reduce NeuroD1 expression when miR-124 level is elevated.

**Figure 1.**
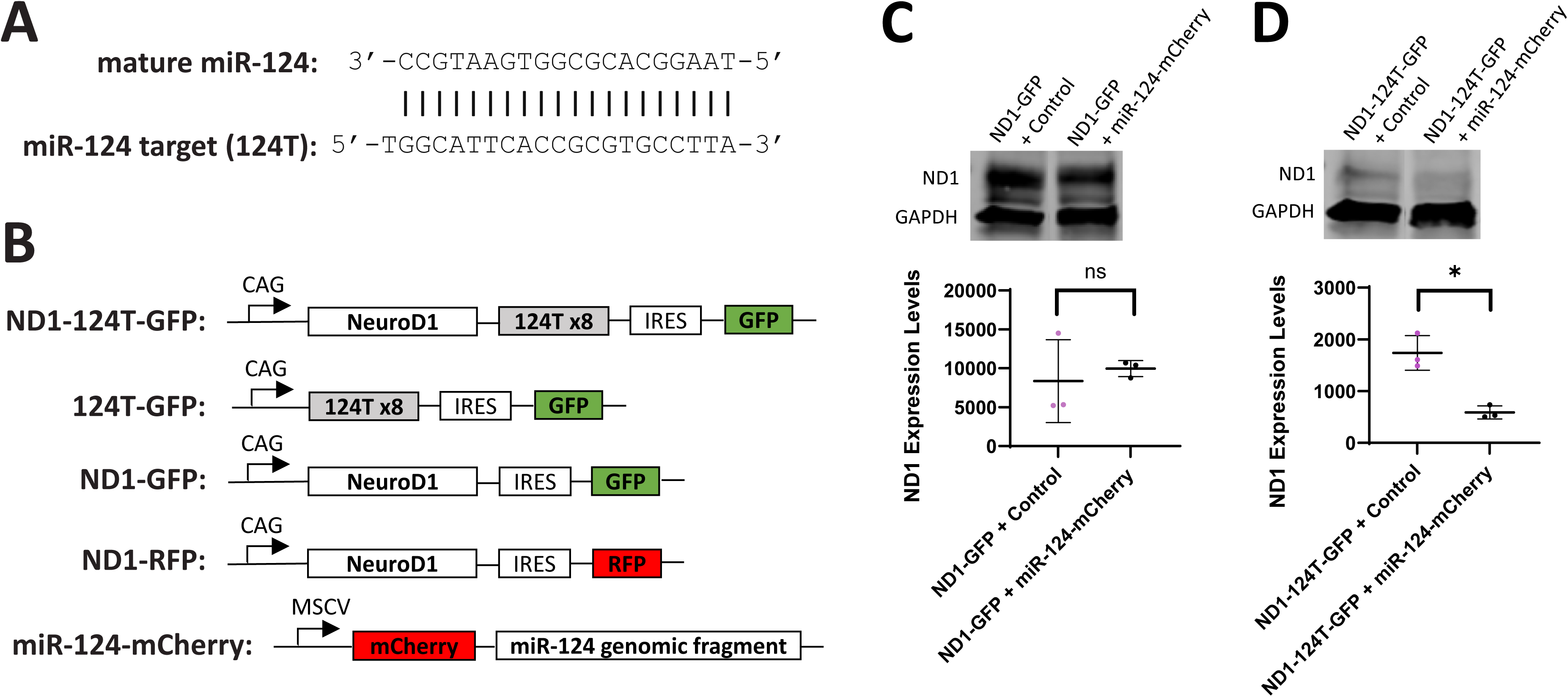
The novel *ND1-124T-GFP* construct is responsive to miR-124 levels. (**A**) Sequence alignment of mature miR-124 and miR-124 target. (**B**) Vector design of the novel *ND1-124T-GFP* construct and other related expression constructs that are used in this study. All vectors contain a strong promoter, either *CAG* or *MSCV*. Western blot analysis was performed on HeLa cell cultures that were co-transfected for 3 days with combinations of either *ND1-GFP* + *miR-124-mCherry* (**C**) or *ND1-124T-GFP* + *miR-124-mCherry* (**D**). NeuroD1 protein levels were normalized by GAPDH and compared between groups. Triplicate samples were included for each group for statistics. ns, not significant; *, p < 0.05 by student *t*-test.

### The ND1-124T-GFP construct elicits a reduced NeuroD1 expression level in HA during neuronal reprogramming

We have previously shown that NeuroD1 induces upregulation of miR-124 in HA as early as 2 days post infection (DPI) during the neuronal reprogramming process^39^. To determine if the *ND1-124T-GFP* construct can respond to the endogenously upregulated miR-124 and modulate NeuroD1 expression level, we first performed western blot analysis on HA cultures that were infected with either *ND1-124T-GFP* or *ND1-GFP* retrovirus. The same titer (10^7^ GC/ml) of the retroviruses were applied for infecting HA cultures, from which cell lysates were prepared at two time points after infection. Our data indicated that the *ND1-124T-GFP* construct elicits a reduced NeuroD1 expression level when compared with *ND1-GFP* at 3 DPI, and this reduction is further enhanced at 6 DPI when the NeuroD1 level by *ND1-124T-GFP* is almost undetectable (**Fig. 2A**). In contrast, the NeuroD1 level by *ND1-GFP* remains constant at these two time points (**Fig. 2A**). We also performed western blot analysis on infected HeLa cells using the same procedure and found no significant reduction in NeuroD1 protein level by *ND1-124T-GFP* at either time points (**Fig. 2B**). Interestingly, HeLa cells express a minimal level of miR-124 and do not upregulate miR-124 expression upon NeuroD1 overexpression (data not shown). Therefore, our western blot analysis indicates that the *ND1-124T-GFP* construct can respond to endogenously upregulated miR-124 and reduce NeuroD1 expression level in a time course manner during NeuroD1-mediated astrocyte-to-neuron reprogramming.

**Figure 2.**
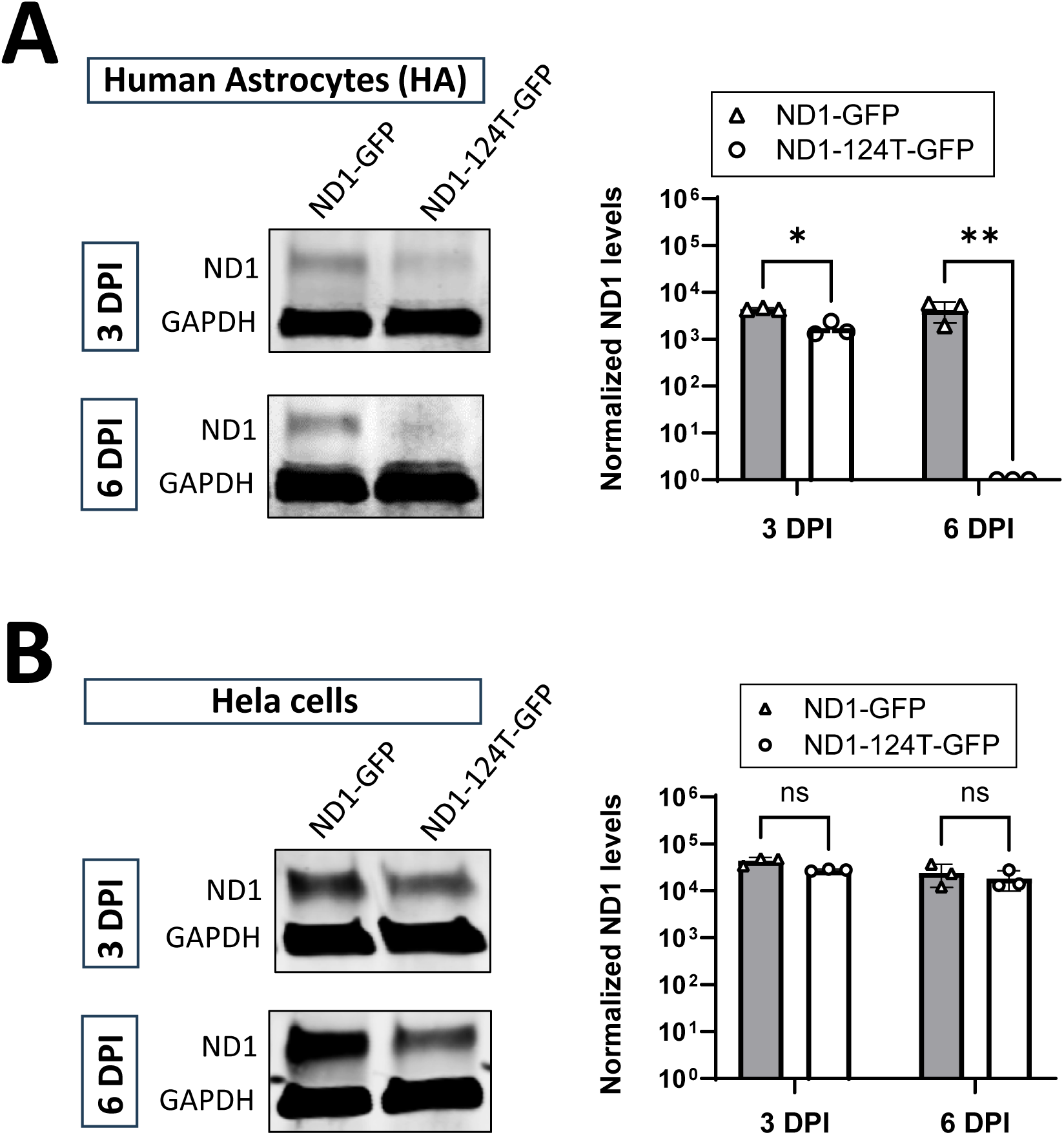
Western blot analysis reveals a decreasing NeuroD1 expression level by the *ND1-124T-GFP* construct in cultured HA but not in HeLa cells. Cultured HA (**A**) and HeLa cells (**B**) were infected with either *ND1-GFP* or *ND1-124T-GFP* retrovirus and harvested at 3- and 6-days post infection (DPI) for Western blot analysis. NeuroD1 protein levels were normalized by GAPDH and compared between groups. Triplicate samples were included for each group for statistics. ns, not significant; *, p < 0.05; **, p < 0.01 by student *t*-test.

To further confirm our western blot results, we measured NeuroD1 expression at single cell level by fluorescence immunocytochemistry using an anti-NeuroD1 antibody. HA cultures were infected with the *ND1-124T-GFP* and *ND1-GFP* retroviruses separately followed by immunostaining. Our results showed that while HA cells infected with *ND1-GFP* exhibit neuronal morphology and strong NeuroD1 levels at 7 DPI (**Fig. 3A**, arrows), the *ND1-124T-GFP* infected HA cells also exhibit neuronal morphology but much reduced level of NeuroD1 (**Fig. 3A**, arrowheads). Statistical analysis indicates that there is a significant reduction of average fluorescence intensity per cell for both NeuroD1 and GFP in the *ND1-124T-GFP* infected cells compared with the *ND1-GFP* infected ones at 7 DPI (**Fig. 3A**). We also performed correlation analysis between cellular GFP and NeuroD1 expression levels in the *ND1-124T-GFP* group and showed that the two levels are both significantly (p<0.0001) and positively (r=0.5198) correlated (**Fig. 3B**). Noting that the fluorescence intensity in immunocytochemistry can be affected by variation factors such as immunostaining procedure and imaging parameters between different coverslips, even though we were cautious in keeping these potential variation factors consistent, we thus designed an infection and mixing experiment (**Fig. 3C**) where we could measure and compare fluorescence intensity of the infected cells on the same coverslips. For that, an ND1 expression construct with an RFP reporter (*ND1-RFP*) (**Fig. 1B**) was included in addition to *ND1-124T-GFP* and *ND1-GFP*. HA cells were infected with the three constructs separately and then passaged and mixed so that each coverslip would have infected cells with two different reporters (**Fig. 3C**). Since *ND1-GFP* and *ND1-RFP* have the same vector design except for different reporters (**Fig. 1B**), they should have the same NeuroD1 expression level, and indeed there is no significant difference in average NeuroD1 fluorescence intensity per cell between *ND1-GFP* (**Fig. 3D**, arrows) and *ND1-RFP* (**Fig. 3D**, asterisks) infected cells at both 3 and 6 DPI (**Fig. 3D**). However, significant differences in NeuroD1 level per cell were detected between *ND1-124T-GFP* (**Fig. 3E**, arrowheads) and *ND1-RFP* (**Fig. 3E**, asterisks) infected cells at both time points with the former being lower (**Fig. 3E**). In addition to reduced NeuroD1 expression levels by *ND1-124T-GFP*, we also observed a diminishing GFP expression of this construct especially at the later time point (**Fig. 3E**, arrowheads). Therefore, our immunocytochemistry analysis further confirms that the *ND1-124T-GFP* construct elicits a reduced level of NeuroD1 expression during astrocyte-to-neuron reprograming in culture.

**Figure 3.**
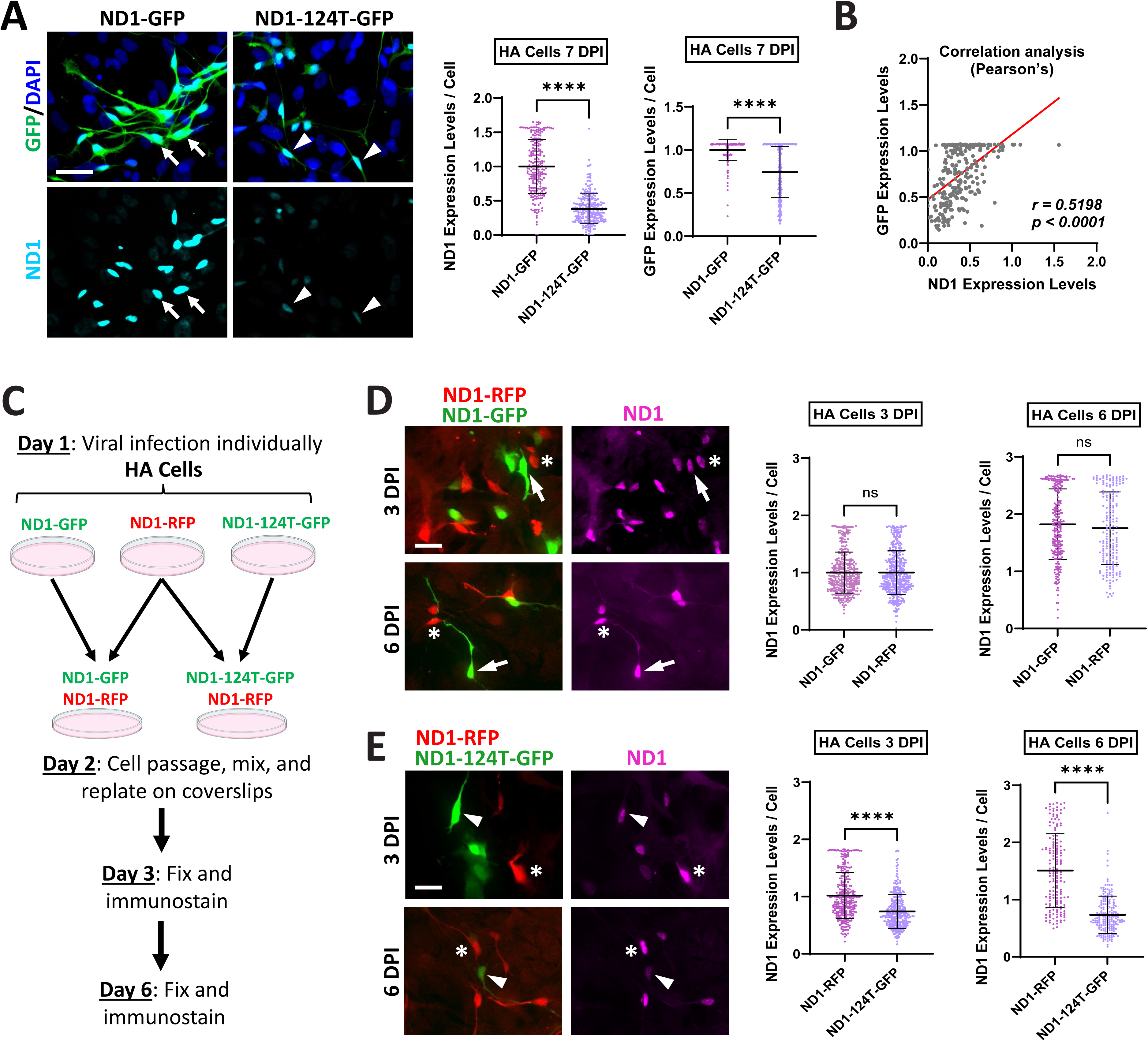
Immunostaining analysis shows reduced NeuroD1 expression levels by the *ND1-124T-GFP* construct in the infected HA during neuronal reprogramming. (**A**) Cultured HA were infected by *ND1-GFP* and *ND1-124T-GFP* retroviruses separately. Following fixation and immunostaining, NeuroD1 protein levels of individual GFP+ cells were measured by fluorescence intensity and compared between the two constructs at 3 and 6 DPI. (B) Correlation analysis (Pearson’s) between GFP and NeuroD1 expression levels among the *ND1-124T-GFP* infected HA cells. (**C**) Diagram depicting the infection and mixing experiment. After fixation and immunostaining, a direct comparison of NeuroD1 expression levels of individual infected cells on the same coverslips was carried out between RFP+ and GFP+ cells. The fluorescence intensity of individual infected cells was measured and compared between *ND1-RFP* and *ND1-GFP* (**D**) and between *ND1-RFP* and *ND1-124T-GFP* (**E**). Arrows, *ND1-GFP* infected cells; *, *ND1-RFP* infected cells; Arrowheads, *ND1-124T-GFP* infected cells in **A**, **D**, **E**. ns, not significant; ****, p < 0.001 by student *t*-test. Scale bars, 40μm.

### The ND1-124T-GFP construct has a dynamically regulated gene expression pattern

According to our original design of the *ND1-124T-GFP* construct, we expect to see a high level of NeuroD1 expression at the early phase during astrocyte-to-neuron reprogramming but reduced NeuroD1 level when miR-124 level is elevated at later stages. Since GFP and NeuroD1 expression levels are positively correlated (**Fig. 3B**), to reveal the dynamic NeuroD1 expression pattern of the *ND1-124T-GFP* construct during reprogramming, we utilized endogenous GFP fluorescence intensity as an indicator of NeuroD1 level in live cells and performed flow cytometry analysis with more time points. HA cultures were infected by both *ND1-124T-GFP* and *ND1-GFP* retroviruses separately and dissociated by trypsinization at different timepoints post infection. After several gating procedures for cell debris and aggregates exclusion, a pool of live single GFP+ cells were subject to detection and readout of GFP intensity (**Fig. 4A**). Our results showed that, while GFP expression level per cell among the *ND1-GFP*-infected HA continues to increase over time, *ND1-124T-GFP*-infected HA reached the peak level of GFP expression at 3 DPI and decreased afterwards (**Fig. 4B**). This dynamic GFP expression pattern of the *ND1-124T-GFP* construct strongly supports our original design that the construct responds to the elevated miR-124 level during NeuroD1-mediated neuronal reprogramming in HA and downregulates transgene expression. We also performed the same experiment in HeLa cells that have a minimal endogenous miR-124 expression and don’t upregulate miR-124 level upon NeuroD1 overexpression as analyzed by qRT-PCR (data not shown). Interestingly, GFP expression level per cell of the *ND1-124T-GFP*-infected HeLa cells lost the up-and-down dynamic pattern as seen in infected HA and continued to increase over time (**Fig. 4C**). On another note, we again observed that GFP expression level by *ND1-GFP* is significantly higher than that by *ND1-124T-GFP* at all the timepoints measured in both HA and HeLa cells (**Fig. 4B and 4C**). Although the exact mechanism of this observation is still unknown, this is likely because the miR-124 target sequences were inserted between the ND1 and GFP coding sequences, which may have affected the efficiency of translation initiation of the internal ribosome entry site (IRES) in the *ND1-124T-GFP* construct (**Fig. 1B**). Nevertheless, even with the suboptimal reporter expression, the *ND1-124T-GFP* construct was successfully demonstrated to have a dynamic transgene expression pattern by flow cytometry analysis during NeuroD1-mediated neuronal reprogramming in cultured HA.

**Figure 4.**
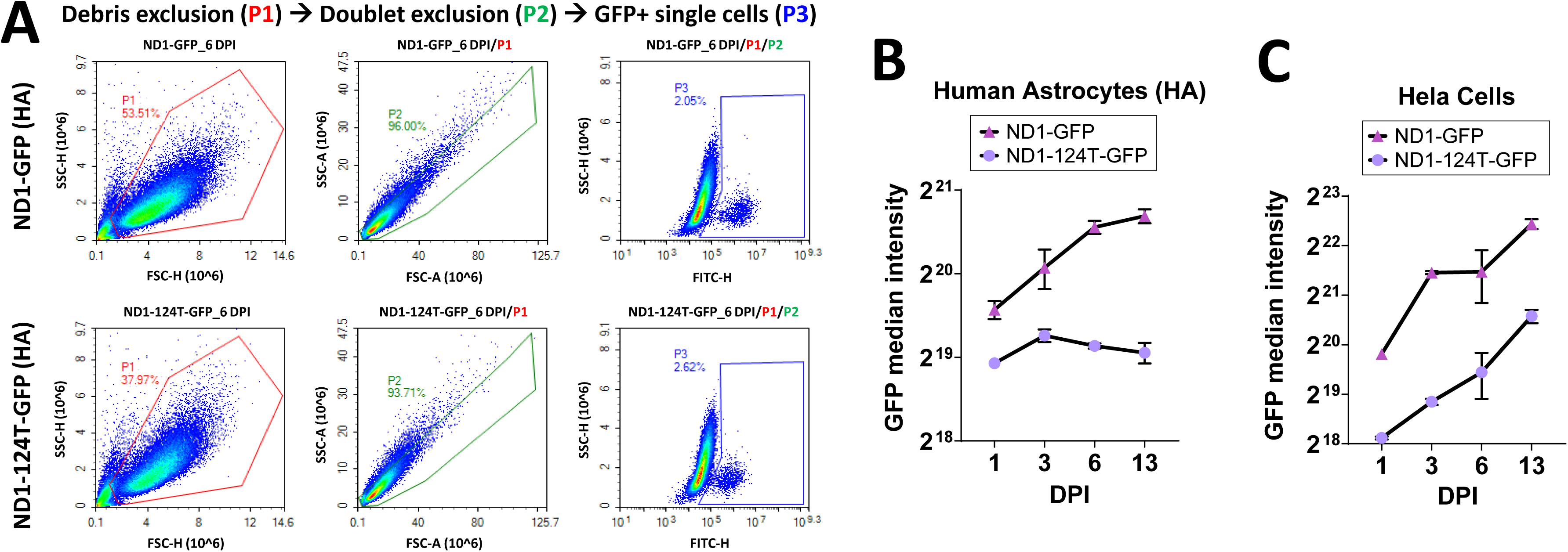
The *ND1-124T-GFP* construct elicits dynamic expression patterns during neuronal reprogramming in HA cultures as revealed by flow cytometry. (**A**) Gating strategy for analyzing individual infected cells (GFP+) by either *ND1-GFP* or *ND1-124T-GFP* constructs. The endogenous GFP expression level of individual live GFP+ cells was quantified at different timepoints, i.e. 1, 3, 6, and 13 DPI, from cultured HA (**B**) and Hela cells (**C**). Triplicate samples were included for each group. The average GFP median intensities of each group were plotted to show time course changes in GFP expression.

### The ND1-124T-GFP construct can efficiently reprogram HA into neurons in culture

After demonstrating the dynamic gene expression pattern of the *ND1-124T-GFP* construct in HA cells, we set out to determine if this new construct possesses neuronal reprogramming capability. We performed retrovirus infection in HA cultures and monitored morphological changes of the infected cells by GFP signal. We found that HA cells infected by either *ND1-GFP* or *ND1-124T-GFP* retrovirus readily change their morphology to elongated neuron-like morphology days after infection. Immunostaining with neuronal markers indicated that these neuron-like cells express both DCX and NeuN at 14 DPI (**Fig. 5A**, arrows). A small number of *ND1-124T-GFP* infected HA cells were often observed to have a flat astrocytic morphology without expression of neuronal markers indicating a lack of neuronal reprogramming (**Fig. 5A**, arrowheads). Quantification analysis showed that the percentage of DCX+ cells did not differ between the two constructs at 7 and 14 DPI, and there is an increase in percentage for both constructs over time reaching nearly 90% at 14 DPI (**Fig. 5B**). On the other hand, a similar increase in the percentage of NeuN+ cells was observed over time for both constructs (**Fig. 5C**). Interestingly, the *ND1-124T-GFP* infected HA showed a slight but significant decrease in this percentage when compared with those of *ND1-GFP* infected HA at 7 DPI, but this decrease becomes insignificant at 14 DPI (**Fig. 5C**). Therefore, these data indicate that *ND1-124T-GFP* is slightly less potent in inducing neuronal reprogramming than *ND1-GFP* but still retains high efficiency.

**Figure 5.**
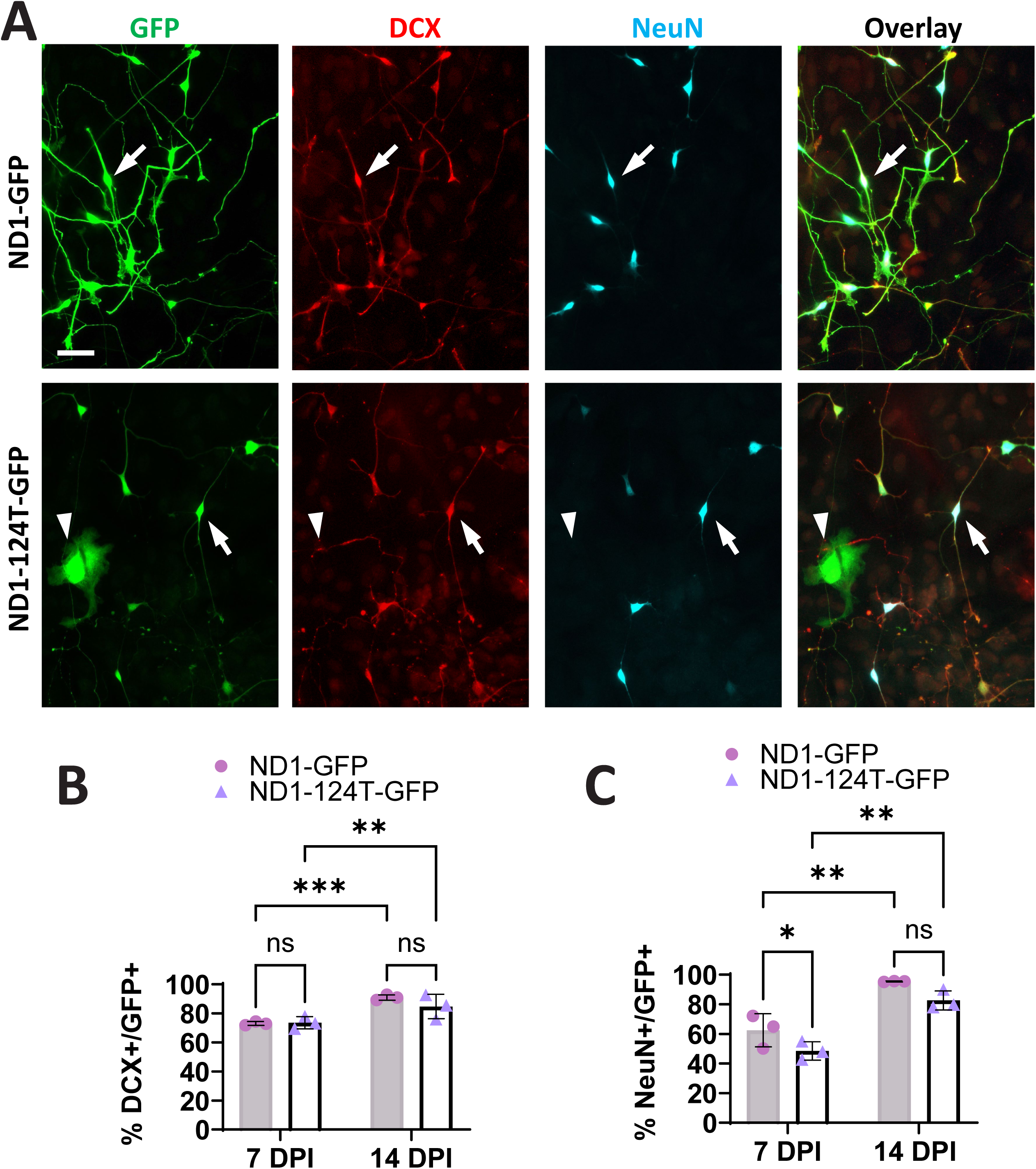
The *ND1-124T-GFP* construct retains relatively high efficiency of astrocyte-to-neuron reprogramming in HA culture. (**A**) Representative images showing immunostaining with neuronal markers DCX and NeuN on HA cells infected by *ND1-GFP* and *ND1-124T-GFP* retroviruses at 14 DPI. Arrows, infected HA cells that are both DCX+ and NeuN+; Arrowhead, a flat cell with neither marker expression that are occasionally seen in the *ND1-124T-GFP* infected HA cultures. Percentages of DCX+ (**B**) and NeuN+ (**C**) cells in the infected HA cultures were quantified and compared at 7 and 14 DPI. ns, not significant; *, p < 0.05; **, p < 0.01; ****, p < 0.001 by student *t*-test. Scale bar, 40μm.

### Overexpression of the miR-124 target sequence itself doesn’t interrupt NeuroD1-mediated neuronal reprogramming

Overexpression of the miR-124 target sequence in the *ND1-124T-GFP* construct may potentially have a “sponge effect”^51^ and reduce cellular level of the mature miR-124 by absorbing it through the complementary sequence. Since miR-124 is not only a marker of mature neurons but also has critical function in neuronal differentiation from neural stem cells^52^, reducing miR-124 level by the sponge effect of miR-124 target sequence may contribute to the observed drop in neuronal reprogramming efficiency of *ND1-124T-GFP* (**Fig. 5C**). To address the question whether the slightly diminished reprogramming potency of *ND1-124T-GFP* is due to the reduced and dynamically regulated NeuroD1 expression level or the potential sponge effect of miR-124 target sequence, we generated a construct expressing only the miR-124 target sequence along with the GFP reporter (*124T-GFP*) (**Fig 1B**). As such, co-infection experiments combining *ND1-RFP* and *124T-GFP* would allow us to specifically test the potential sponge effect of miR-124 target sequence in astrocyte-to-neuron reprogramming. We first validated the specificity of the *124T-GFP* construct by co-infection with overexpression vectors of either miR-124 or miR-375 (an unrelated miRNA that is also induced during neuronal reprogramming^39^) that carry a mCherry reporter. Interestingly, double-positive cells strongly expressing mCherry and GFP were often observed in HA cultures that were co-infected by *124T-GFP* and *miR-375-mCherry* at 3 DPI (**Fig. 6A**, arrows), but rarely observed in *124T-GFP* and *miR-124-mCherry* co-infected cultures (**Fig. 6A**). This result indicates that the *124T-GFP* construct responds specifically to miR-124 level by reducing reporter expression (like a miR-124 sensor) but not to miR-375, and it also suggests its specificity of the potential sponge effect. We then co-infected HA cells with *ND1-RFP* and *124T-GFP* retroviruses. The co-infected cells (**Fig. 6B**, arrows) were readily observed at 3 DPI and changed to neuronal morphology with long processes at 7 DPI mimicking the HA cells infected by *ND1-RFP* retrovirus only (**Fig. 6B**, arrowheads). Furthermore, the co-infected HA cells acquired enhanced neuronal morphology and expressed neuronal markers DCX and NeuN at 14 DPI (**Fig. 6C**, arrows). We also noticed a much weaker GFP signal of the co-infected cells (**Fig. 6C**, arrows) compared with the *124T-GFP* only cells exhibiting a flat morphology in the same cultures, which is consistent with the proposed miR-124 sensor activity of this construct (**Fig. 6A**). Quantification analysis showed that no significant difference in the percentages of marker+ cells was observed when comparing co-infected HA with either *ND1-RFP* singly infected HA in the same co-infection experiment (**Fig. 6C**, arrowheads) or HA cells infected by *ND1-RFP* only in a separate experiment (**Fig. 6D**, arrowheads). Taken together, these results suggest that the slightly compromised neuronal reprogramming efficiency of the *ND1-124T-GFP* construct is not due to the potential sponge effect of miR-124 target sequence but rather its reduced NeuroD1 expression level.

**Figure 6.**
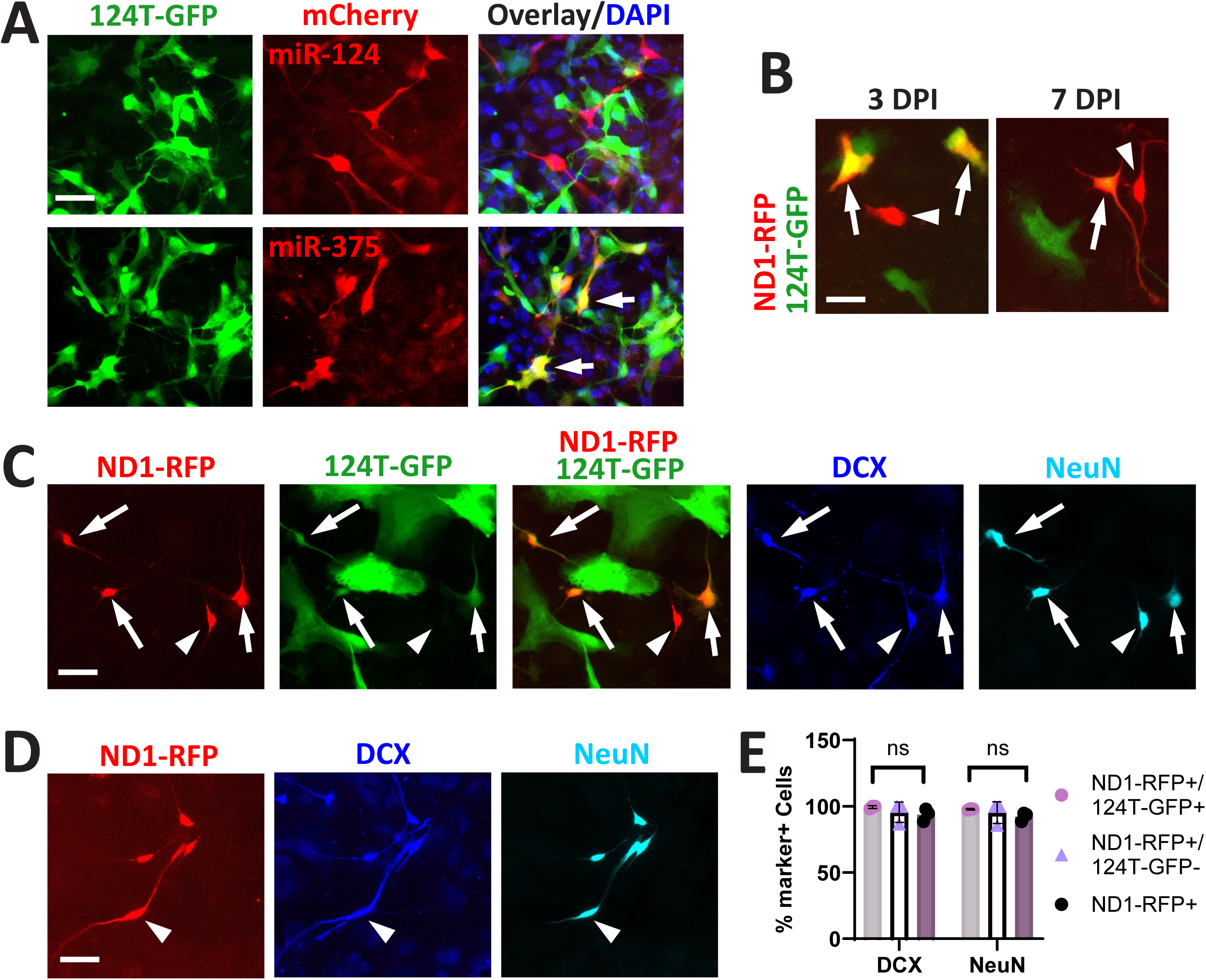
Overexpression of the miR-124 target sequence itself does not affect NeuroD1-mediated astrocyte-to-neuron reprogramming in HA culture. (**A**) Live fluorescence images of GFP and mCherry in HA cultures co-infected by *124T-GFP* retrovirus with either *miR-124-mCherry* or *miR-375-mCherry* retroviruses at 3 DPI. Arrows, co-infected HA cells with strong GFP and mCherry expression. (**B**) Live fluorescence images of GFP and RFP in HA cultures co-infected by *124T-GFP* and *ND1-RFP* retroviruses at 3 and 7 DPI. Immunostainings with antibodies against neuronal markers DCX and NeuN were performed on HA cultures either co-infected by *ND1-RFP* and *124T-GFP* retroviruses (**C**) or infected by *ND1-RFP* only (**D**) at 14 DPI. (**E**) Quantification and comparison of percentages of marker+ cells in **C** and **D**. Arrows, co-infected HA cells; Arrowheads, HA cells infected by *ND1-RFP* only. ns, not significant by student *t*-test. Scale bars, 40μm.

### The reprogrammed neurons by ND1-124T-GFP can mature and express neuronal subtype markers in long term cultures

Next, we set out to determine if the *ND1-124T-GFP*-reprogrammed neurons can survive in long term cultures and express markers of mature neurons and neuronal subtypes. To promote cell survival and neuronal maturation, we adopted a 3-D spheroid culture system to increase cellular density and mimic in vivo situations. Spheroids of HA cells were generated at 3 DPI following a modified hanging drop protocol^50^ (**Fig. 7A**) and replated onto a monolayer of HA cells on the next day. The spheroids of infected HA cells were monitored over time on their reprogramming process. At later stages, the reprogrammed neurons formed clusters with intermingled neuronal processes in the core (**Fig. 7B**). Some reprogrammed neurons migrated out to the periphery and exhibited a typical neuronal morphology (**Fig. 7B**, arrowhead). The spheroid cultures were usually maintained for 30 days with proper medium change before being fixed. Immunostaining analysis showed that the reprogrammed neurons by *ND1-124T-GFP* express mature neuronal markers (NeuN and Map2) and exhibit nice neuronal morphology similarly to the ones by *ND1-GFP* (**Fig. 7C**). Interestingly, while the *ND1-GFP*-reprogrammed neurons still express high level of NeuroD1 in their nuclei, the *ND1-124T-GFP*-reprogrammed ones have a much lower level by comparison (**Fig. 7C**, arrows) indicating that the reduced NeuroD1 level by *ND1-124T-GFP* can be maintained long term in the reprogrammed neurons. We also detected expression of the synaptic vesicle marker SV2 in both cultures (**Fig. 7D**, arrows). For neuronal subtype identification, our immunostaining results showed that reprogrammed neurons by both constructs express markers of glutamatergic neurons vGlut1, HuD (**Fig. 7E**), and Ctip2 (**Fig. 7F**) at this stage. However, the expression levels of these markers are mostly decreased in the *ND1-124T-GFP*-reprogrammed neurons (**Fig. 7G**). Surprisingly, both reprogrammed neurons also express GABAergic neuronal markers GABA and GAD67 (**Fig. 7H**), albeit the *ND1-124T-GFP*-reprogrammed neurons have a more enhanced expression level of these markers by comparison (**Fig. 7I**). Taken together, these results indicate that the reprogrammed neurons by *ND1-124T-GFP* can be mature in long term cultures and elicit reduced glutamatergic neuron marker expression and enhanced GABAergic neuron markers compared with the *ND1-GFP*-reprogrammed neurons.

**Figure 7.**
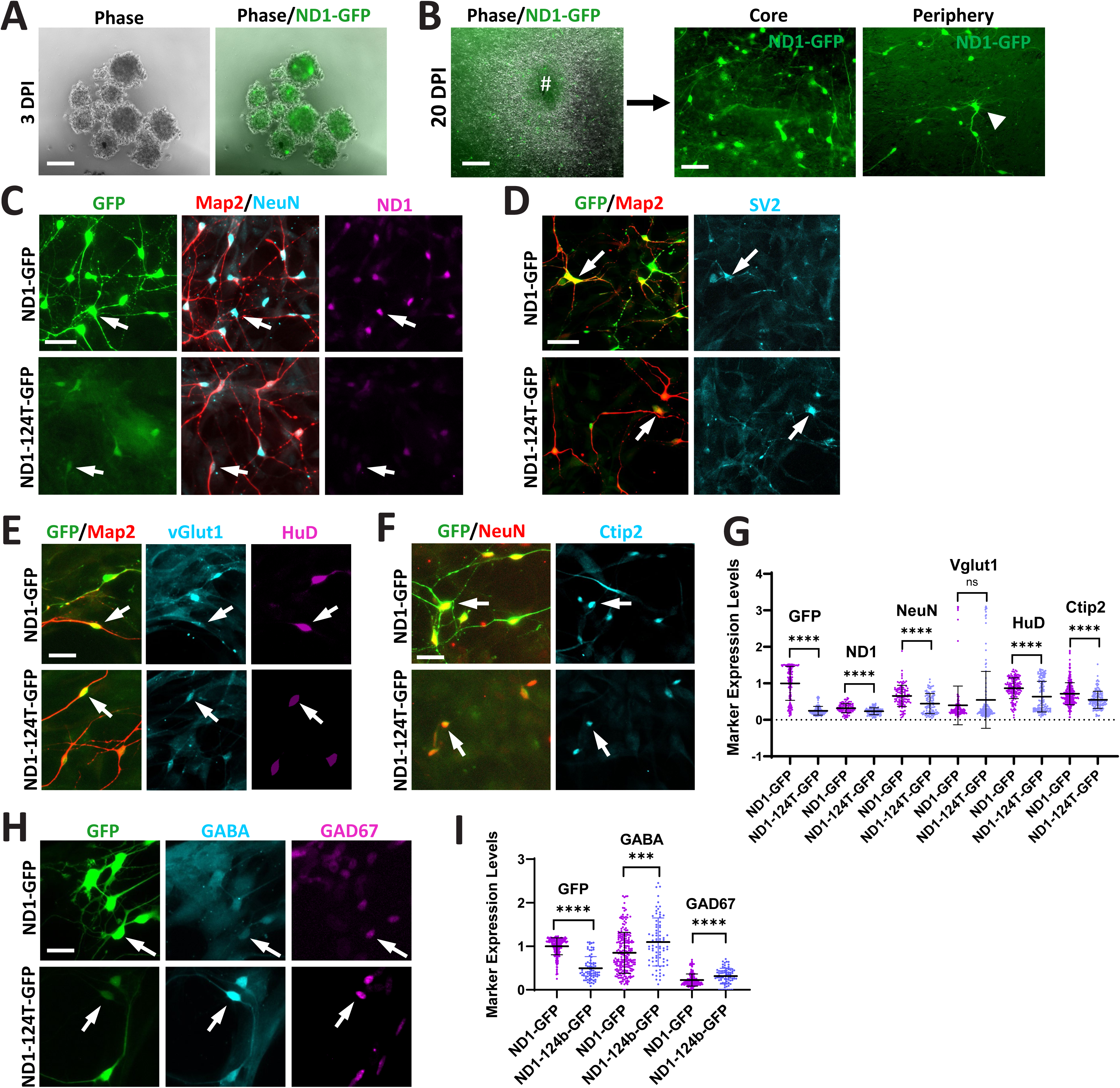
The reprogrammed neurons by the *ND1-124T-GFP* construct can express markers of mature neurons and neuronal subtypes in long term cultures. (**A**) A live image of HA cell cultures infected by *ND1-GFP* retrovirus showing floating cell spheroids in the hanging drops of culture medium at 3 DPI. (**B**) Live images of the resulting cell spheroids that are attached to a monolayer of HA cells at 20 DPI. #, the core of the attached spheroid; Arrowhead, a HA cell infected by *ND1-GFP* that has migrated to periphery of the spheroid showing a typical neuronal morphology. At 30 DPI, the attached HA spheroid cultures infected by either *ND1-GFP* or *ND1-124T-GFP* were fixed and immunostained with the indicated mature neuronal markers (NeuN and/or Map2) along with NeuroD1 (**C**), the synaptic vesicle marker SV2 (**D**), the glutamatergic neuronal markers vGlut1 and HuD (**E**), and Ctip2 (**F**). (**G**) Expression levels of the markers were quantified by their cellular fluorescence intensities and compared. (**H**) The GABAergic interneuron markers GABA and GAD67 (**F**) were also analyzed and compared (**I**). Arrows, the marker+ cells. ns, not significant; ***, p < 0.005; ****, p < 0.001 by student *t*-test. Scale bars, 40μm.

## Discussion

In this study, we engineered a novel expression construct and tested the proof of principle if a dynamic gene expression pattern can be achieved in the model of astrocyte-to-neuron reprogramming. The uniqueness of this construct is the incorporation of a miRNA responsive element by which transgene expression can be modulated when target cells change their cellular identity from astrocytes to neurons during the reprogramming process. Selecting miRNAs for this design is crucial. We wanted to select the miRNAs that have low levels in astrocytes but increased levels in reprogrammed neurons so that, when combined with a strong promoter, the transgene expression can elicit a dynamic pattern, i.e. from high to low, resulting from the inhibitory mechanism of these miRNAs. The expression pattern of miR-124 meets the requirement for this design, and therefore it is selected for this study although there might be other candidate miRNAs that can be tested in the future. In vivo neuronal reprogramming has emerged as a promising alternative strategy for regenerative medicine. Having been demonstrated for feasibility in numerous disease/injury models^2,3,53–55^, the strategy would require optimization for moving toward clinical applications. Unlike cell reprogramming in vitro where exogenous factors and drugs can be easily supplemented in culture to manipulate the reprogramming process, it is challenging to do so in vivo although not impossible^56,57^. Therefore, the outcome of in vivo reprogramming largely depends on the intrinsic properties of the expression construct of the reprogramming factors, and optimization of the vector design for a successful reprogramming outcome^13^ would become a very important research area in this regard. Here, we provide experimental data to show that the *ND1-124T-GFP* construct expresses the reprogramming factor NeuroD1 in a dynamic fashion and results in a much-reduced NeuroD1 level in the reprogrammed neurons, which may alter their neuronal subtype identity.

One exciting piece of data from this study is the demonstration of a dynamic expression pattern of the *ND1-124T-GFP* construct during astrocyte-to-neuron reprogramming process by flow cytometry (**Fig. 4**). This data confirmed that our expression construct design is working as expected in the model of reprogramming. Even though the flow cytometry monitors the time-course expression of GFP, we reason that NeuroD1 expression would follow a similar pattern since both transgenes are on the same vector. However, GFP and NeuroD1 proteins may have different turnover rates, which will affect the final concentrations of these proteins in the cell. In fact, NeuroD1, as a transcriptional factor, undergoes rapid protein degradation through the ubiquitin-dependent proteasomal pathway^58^, while the GFP reporter normally has a half-life as long as 26 hours in the cell^59^. Therefore, we expect that NeuroD1 would have a similar dynamic expression pattern to GFP during the reprogramming process but probably a much quicker decline at the protein level than GFP in the reprogrammed neurons. Reliably detecting NeuroD1 protein levels by flow cytometry has failed in our hand after testing several anti-NeuroD1 antibodies. Nevertheless, we were able to show the decreased level of NeuroD1 protein by immunocytochemistry in the *ND1-124T-GFP*-reprogrammed neurons (**Fig. 3**), which is consistent with the pattern of GFP expression as demonstrated by flow cytometry. The expression level and duration of the reprogramming factors is important for a successful reprogramming outcome as discussed in our recent review^13^. Although transient NeuroD1 expression is sufficient to induce neuronal reprogramming from cultured fibroblasts^60^, in another example, NeuroD1 expression must reach a threshold level to convert microglia to neurons in culture^12^. Given that *ND1-124T-GFP* can induce neuronal reprogramming with relatively high efficiency (**Fig. 5**), its NeuroD1 expression level should have reached a sufficient level, and the subsequent lower level would not affect reprogramming per se. In fact, it is this lower level of NeuroD1 after reprogramming that we think could affect the behavior of the resulting neurons such as their neuronal subtype identity.

We performed an experiment to address the potential sponge effect of miR-124 target sequence and showed that overexpression of *124T-GFP* during NeuroD1-mediated neuronal reprogramming does not affect reprogramming efficiency as measured by percentages of DCX+ and NeuN+ cells (**Fig. 6**). However, whether it has adverse effects in the long term such as neuronal maturation and function is still unclear and needs to be addressed in future studies. Gene knockout studies have shown that miR-124 is not essential to neurogenesis but may affect physiological functions such as synaptic formation when deleted^61,62^. We don’t know to what degree the sponge effect of the miR-124 target sequence would affect the total concentration of cellular miR-124 in the reprogrammed neurons. Also, we don’t know if this potentially reduced miR-124 level would have a significant effect on gene expression and thus changes in neuronal behavior. A thorough characterization of the reprogrammed neurons in long-term cultures and in vivo settings would be needed to answer these questions. If the sponge effect becomes a problem in the reprogrammed neurons, an alternative approach is to use other miRNAs that are less crucial than miR-124. Our previous report demonstrated that miR-375 is induced drastically by NeuroD1 overexpression in HA^39^ and could be an alternative candidate.

Being able to generate reprogrammed neurons with distinct neuronal subtypes, i.e. excitatory vs. inhibitory, will increase the flexibility of the reprogramming approach for regenerative medicine. Our hypothesis is that continuously high level of NeuroD1 drives the reprogrammed neurons to glutamatergic phenotype, and that by reducing NeuroD1 level after the reprogramming process will diminish this driving force and allow acquisition of other subtype identities. The neuronal subtype identity of the *ND1-124T-GFP*-reprogrammed neurons were characterized in a long-term culture condition. Upon initial assessment, we observed many reprogrammed neurons co-express markers of GABAergic and glutamatergic neuronal subtypes (data not shown). To unbiasedly analyze neuronal subtypes, instead of calling marker+ neurons and calculating percentages, we decided to measure expression levels of the markers in every infected cell for statistical comparison. Indeed, the *ND1-124T-GFP*-reprogrammed neurons showed higher expression levels of GABAergic neuron markers and lower levels of glutamatergic neuron markers by comparison (**Fig. 7**). Co-expression of markers of both neuronal subtypes could be a culture artifact since expression of these markers can be modulated by exogenous factors. For example, BDNF, PDGF and SHH have been shown to promote the GABAergic neuron phenotype in cultures^63,64^. On the other hand, a dual GABAergic/glutamatergic neuronal phenotype has been identified in numerous regions of the CNS^65–67^. It could be that the reduced level of NeuroD1 in the *ND1-124T-GFP*-reprogrammed neurons makes them prone to become other neuronal subtypes, and additional environmental factors may drive them to one subtype over the other. Therefore, in vivo settings may be a good condition for testing the subtype identification of these reprogrammed neurons and will be applied in our future research. In consistent with our hypothesis, NeuroD1 with different expression levels reprograms astrocytes to different subtypes of neurons in the mouse retina^68^.

In conclusion, our current study has characterized a novel expression construct *ND1-124T-GFP* and showed that by incorporating a miR-124 target sequence, it exhibits a dynamic expression pattern of NeuroD1 during the astrocyte-to-neuron reprogramming process and could be used to generate new neurons with diversified neuronal subtypes for a wide range of regenerative applications.

## Author contributions

H.L. and X.C. conceived the idea and supervised the entire project. N.J. performed the major experiments, analyzed the data, and made the figures. M.J., M.S., A.R., and C.W. contributed to the experiments. H.L. wrote the manuscript and secured funding.

## Data and materials availability

The raw data supporting the conclusions of this article will be made available by the authors, without undue reservation.

## Acknowledgements

This work was supported by startup funds from Medical College of Georgia at Augusta University, National Institutes of Health grants (R01NS117918, R21NS104394, R21NS119732), and Ann L. Jones Spinal Cord Regeneration Research Fund.

